# Rapid Molecular Mechanotyping with Microfluidic Force Spectroscopy

**DOI:** 10.1101/2023.02.17.528971

**Authors:** Martijn van Galen, Annemarie Bok, Taieesa Peshkovsky, Jasper van der Gucht, Bauke Albada, Joris Sprakel

## Abstract

Molecular mechanotyping, the quantification of changes in the stability of supramolecular interactions and chemical bonds under the action of mechanical forces, is an essential tool in the field of mechanochemistry. This is conventionally done in single-molecule force-spectroscopy (smFS) assays, for example with optical tweezers or Atomic Force Microscopy. While these techniques provide detailed mechanochemical insights, they are time-consuming, technically demanding and expensive; as a result, high-throughput screening of the mechanochemical properties of molecules of interest is challenging. To resolve this, we present a rapid, simple and low-cost mechanotyping assay: microfluidic force spectroscopy (*µ*FFS), which probes force-dependent bond stability by measuring the detachment of microparticles, bound to microfluidic channels by the interaction of interest, under hydrodynamic forcing. As this allows the simultaneous observation of hundreds of microparticles, we obtain a quantitative mechanotype in a single measurement, using readily available equipment. We validate our method by studying the stability of DNA duplexes, previously characterized through smFS. We further show that we can quantitatively describe the experimental data with simulations, which allows us to link the *µ*FFS data to single-bond mechanochemical properties. This opens the way to use (*µ*FFS) as a rapid molecular mechanotyping tool.

## Introduction

Understanding how the stability, e.g. the dissociation rate, of a supramolecular interaction or chemical bond changes under the action of mechanical forces is a crucial step in a variety of topics in both mechanochemistry and mechanobiology. Examples include the development of mechanically-activated chemical reactivity to strengthen materials, ^1,2^ the mechanical gating of catalytic activity in both synthetic materials and nature,^3,4^ or mechanotransduction pathways in cells by means of proteinaceous catch bonds.^5–7^ The susceptibility of bonds to mechanical dissociation is typically evaluated by measuring the lifetime of the interaction, i.e. the inverse of the bond dissociation rate, under the action of tensile forces. For many bonds, including covalent chemical bonds and supramolecular interactions, the lifetime decreases exponentially with force, as described by the Bell-equation:^8^

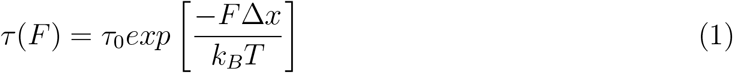

where *τ* is the bond lifetime, *τ*_0_ its value in the absence of force, *F* the tensile force on the bond and Δ*x* the so-called activation length, a property that sets the susceptibility of a bond to mechanical rupture. For bonds that obey Bell-type mechanochemistry, conventionally known as slip bonds, mechanotyping comes down to measuring the force-lifetime *τ* (*F*) curve, so that the parameters *τ*_0_ and Δ*x* can be extracted.

Force-lifetime curves are usually obtained through force-clamp measurements by means of single-molecule force spectroscopy (smFS) techniques. These techniques use single-molecule manipulation, e.g. in optical tweezers or an Atomic Force Microscope, to place individual bonds under tension and record their dissociation kinetics. ^9^ Mechanotyping involves numerous repeats of these experiments over a range of tensile forces.^10^ In the past decades, smFS has become an invaluable method to study mechanochemistry at the level of single molecules, having provided critical insights into a wide variety of phenomena, ranging from chemical bonds^11^ and synthetic supramolecular interactions,^12^ to the stability and folding trajectories of proteins,^13^ and to the stability of DNA or even entire chromosomes.^14,15^ Despite their success, current smFS methods can be challenging. They are sophisticated techniques that require expensive equipment and extensive user experience to operate,^10^ making it difficult to perform these experiments in a typical mechanochemistry or mechanobiology laboratory. Moreover, the measurement principle of probing one molecule at a time poses a challenge in itself: As single bond rupture events are stochastic by nature, every measurement must be repeated a large number of times to obtain sufficient statistics to assess the most-probable bond lifetime for any given applied force. This makes these methods time-consuming.

Over time, several smFS techniques have resolved these challenges, by employing clever parallelization schemes, such as those used in magnetic tweezers or acoustic force spectroscopy.^16–18^ These techniques can make many measurements simultaneously, by binding a large number of microparticles, through the interaction of interest, to the surface of a microfluidic chip. An equal amount of tension is then applied to all particles simultaneously, either by magnetic or by acoustic excitation, and their detachment is monitored using a simple microscope. While such parallelization schemes substantially speed up molecular mechanotyping, they still require substantial investments in specialised instrumentation.

Here we set out to implement the principle of parallelized mechanotyping by applying forces in a simple and cheap microfluidic device, making use of the well-defined hydrodynamic forces that the laminar flow inside the chips provides. We present MicroFluidic Force Spectroscopy (*µ*FFS), a rapid mechanotyping assay that allows the measurement of the mechanotype of (bio)molecular interactions in a single afternoon, with sufficient datapoints to enable a robust determination of the molecular mechanotype. We explore our method by studying the stability of DNA duplexes, and find excellent agreement with previous results on the same interactions by smFS. We also develop a simulation model, that not only quantitatively maps onto the experimental data, but provides insight into the molecular details of the experiments.

This opens the way to use a combination of *µ*FFS experiments and simulations as a rapid mechanochemical screening tool.

## Results and discussion

Our *µ*FFS approach allows us to measure the strength of (bio)molecular interactions using only a disposable microfluidic chip, a flow-pressure controller connected to a flow-rate sensor, and a brightfield microscope (Fig. 1A). First, the flow-pressure controller (1) pumps a dispersion of polystyrene microparticles, fuctionalised with one of the two interaction partners (2) into a PDMS microfluidics chip, whose bottom glass substrate is functionalised with the complementary interaction partner (3), which is mounted on the brightfield microscope. After passing the chip, the fluid is passed through a flow rate sensor (4) that allows for accurate flow rate control through a feedback loop flow controller. A detailed description is given in the Materials and Methods section.

**Figure 1:**
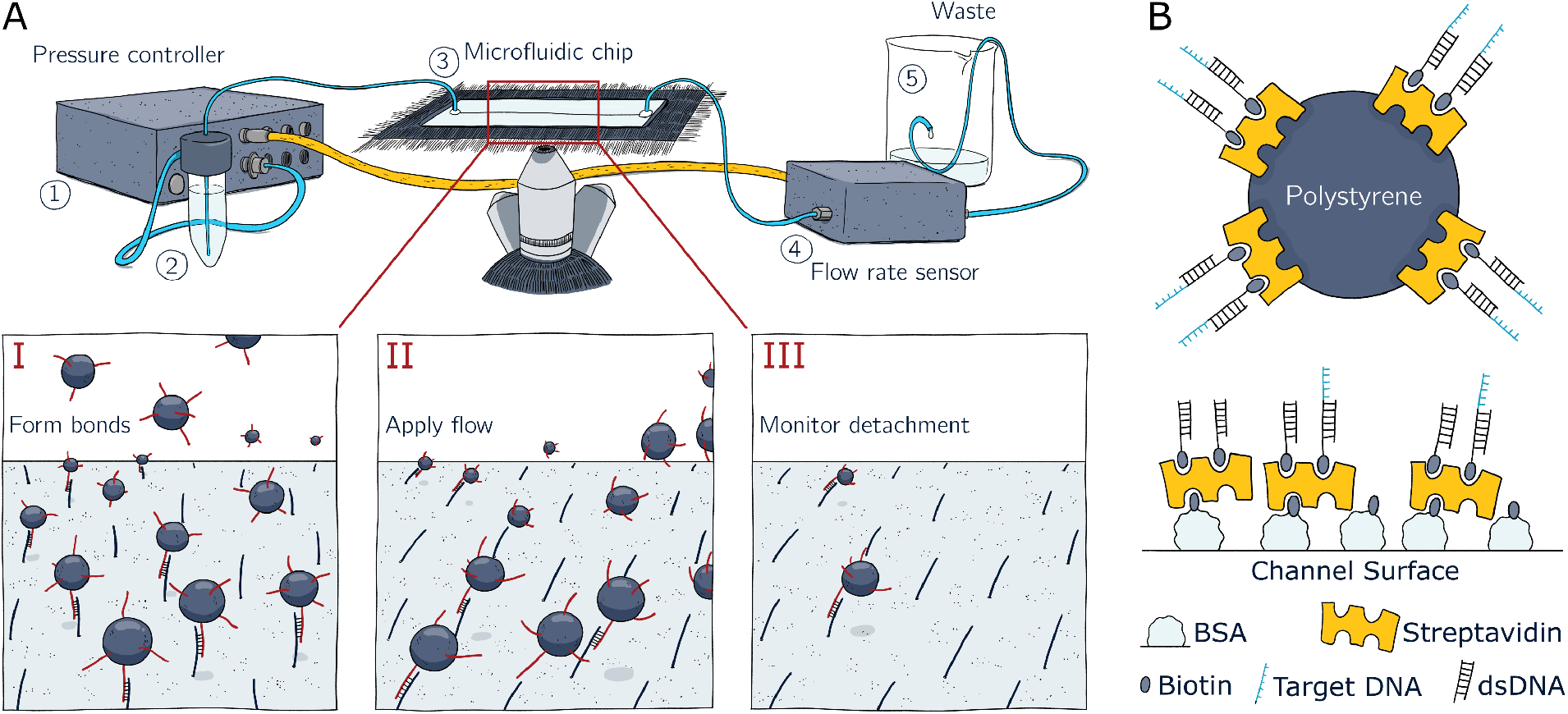
A: Schematic representation of the microfluidic force spectroscopy setup. A pressure controller (1) is used to pump a suspension of polystyrene particles (2) through a microfluidics chip (3) mounted on a brightfield microscope. A flow rate sensor (4) is used to maintain a set flowrate through a feedback loop with the pressure controller. The remaining flow-through is disposed in (5) a waste container. A measurement consists of (I) allowing particles to settle and form bonds with the surface, (II) switching on the flow rate to apply a shear stress on the particles, followed by (III) monitoring particle detachment over time. B: Surface chemistry used to attach the target DNA bond to the particles and microfluidics channels. Top: Streptavidin-coated polystyrene beads are functionalized with biotinylated DNA. Bottom: Channel surfaces are first coated in biotinylated Bovine Serum Albumin, followed by streptavidin and finally by biotinylated DNA. A Mixture of target DNA and nonspecific DNA is used to tune the contact valency.

A *µ*FFS experiment is performed as follows: After functionalisation of the particles and chip with the interaction partners of interest, (I): the dispersion of particles is introduced into the chip and the flow is switched off. As the particles are slightly denser than their suspending medium, they sediment and form adhesive bonds with the surface of the chip. (II): After five minutes of incubation, the flow is switched on, with a short ramp-up time of *<*1 second, to a target flow rate *Q*. The fixed applied flux *Q* results in a well-defined shear stress at the bottom surface, resulting in a torque *M* and a shear force *F*_*shear*_ on the particles.^19,20^ This constitutes the equivalent of a force-clamp experiment in smFS, where a force is rapidly applied to the interaction of interest and kept constant, waiting for bond dissociation to occur.

As the adhesive interactions between particle and substrate dissociate under the applied force, particles detach from the surface. (III) By recording videos of the particles using the microscope, and by particle tracking analysis, we quantify particle detachment kinetics, and construct bond survival probability histograms, from which we can determine the characteristic bond lifetime *τ*. By repeating these measurements as a function of the applied flux *Q*, i.e. by varying the applied force *F*_*shear*_, we can construct a force-lifetime curve *τ* (*F*_*shear*_).

To validate the *µ*FFS method, we study interactions whose mechanochemical properties are known from extensive smFS measurements reported in literature.^21^ We use short DNA duplexes with sizes ranging between 15-30 nucleotides (Table 2). Performing the measurements requires a strategy to tether the interaction partners to the beads and chip surface, which ideally is versatile and simple. One strategy that meets these requirements is the use of biotin-streptavidin interactions (Fig. 1B). These interactions rely on the bonding between the cofactor biotin in one of the four binding pockets of the streptavidin protein; forming one of the strongest natural supramolecular interactions, with reported rupture forces in the range of 100 to 300 pN. ^22^ As such, this approach is only suitable for measuring bio(molecular) interactions that are substantially weaker than the biotin-streptavidin interaction, although it could be extended to stronger bonds by using covalent functionalisation strategies.

We first functionalize the bottom surface of the microfluidic channels with streptavidin by physisorption of a biotinylated bovine serum albumin (BSA) protein, followed by incubation with streptavidin, and we use commercial streptavidin-coated PS beads. Our DNA duplexes consist of a stem: a 40bp DNA duplex, and a reactive overhang that is available for bonding. As we want control over the bonding valency in the adhesive contact, we require a means to vary the surface density. We therefore functionalize the microfluidic channels with a mixture of DNA oligomers containing a sticky end that is complementary to the DNA on the particles (=target DNA) and DNA oligos containing only the 40 bp duplex stem (=filler DNA). By varying the molar ratio of target and filler DNA, we can vary the surface density of reactive bonds. All DNA sequences used in this study are shown in Table 2 in the Materials and Methods section.

We first test whether our functionalization strategy provides a homogeneous distribution of DNA around the particles available to bind its conjugates. For this, we incubated particles functionalized with the target DNA with a complementary DNA strand that carries a Alexa488 fluorescent reporter (Ftest-A488, Table 2). Confocal microscopy images show homogeneous functionalisation, and only slight variations in fluorescence intensity between different particles, which is likely due to a variation in streptavidin loading between the particles (Fig. 2A). Using a pull-down assay, we quantified the concentration of available DNA linkers on the particle to be 3.9 · 10^3^ sites*/µ*m^2^ (SI, section S4). We performed a similar fluorescence experiment on the surface of the microfluidic chip. Upon increasing the ratio of target DNA to inert filler DNA, we find a clear increase in the fluorescence intensity, signalling control over the density of available binding sites (Fig. 2B). The chip functionalization also appears homogeneous, although minor patchiness can be observed around 50% of target DNA. Interestingly, quantification of the fluorescence intensity as a function of the ratio of reactive to inert filler DNA reveals a non-linear relationship between the fluorescence intensity and the fraction of target DNA, suggesting that the fraction of target DNA in solution does not correspond linearly to the fraction of target DNA at the surface. This is most likely due to differences in binding affinity to streptavidin between the target DNA and filler DNA caused by their differences in size and rigidity.

**Figure 2:**
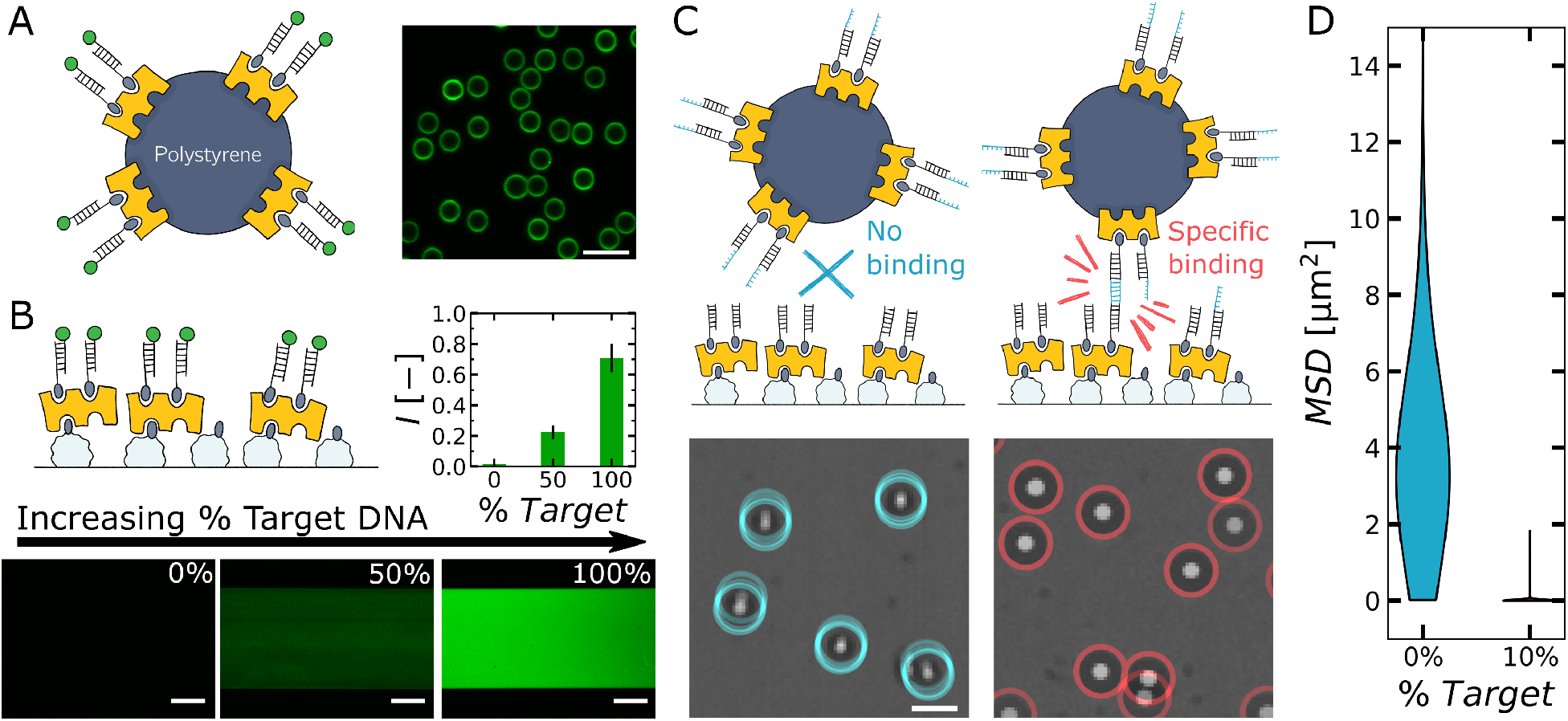
Fluorescence tests with an Alexa488-labeled DNA conjugate show homogeneously functionalized surfaces on (A) the particles and (B) the microfluidics chip. B: Increasing the fraction of target DNA on the surfaces results in an increase of the fluorescence intensity. C: Binding specificity test to confirm that particles adhere to the channel surface through DNA base pairing interactions. Particles remain Brownian on surfaces with 0% target DNA (left), but firmly adhere to surfaces with 10% target DNA (right). We quantify this effect by computing the mean-square displacement (MSD) of each particle over five seconds (D) and show that the two surfaces are significantly different. Scale bars denote A: 10 *µ*m, B: 500 *µ*m, and C: 10 *µ*m.

All molecular mechanotyping experiments can be troubled by the emergence of non-specific interactions between particle and surface that can preclude accurate measurement of the properties of the specific interaction of interest, and which are difficult to filter out post-measurement. To assess this, we performed a binding specificity test in which we functionalized the surface with 0% target DNA and 100% filler DNA, so that there should be no specific interaction between particles and surface. The absence of non-specific bonding was verified by the observation that particles in contact with a non-reactive DNA coated chip remained fully mobile after sedimentation; showing diffusive transport over multiple micrometers during five second time-lapse movies (Fig.2C). By contrast, the presence of even small amounts of reactive target DNA resulted in immobile particles after sedimentation. This effect was further quantified by computing a histogram of the mean-squared displacement of the particles over a five second period, which is much larger on the 0% target DNA surface than it is on surfaces functionalised with 10% target DNA (Fig.2D). This shows that our functionalisation procedure results in surfaces that are free from non-specific interactions, and hence that any mechanically-stable interactions we observe must be due to DNA base pairing.

Having established the binding specificity, we now turn to mechanotyping of DNA duplexes. These experiments were performed as follows: First, we introduce the functionalized particles into the functionalized devices, stop the flow and begin recording time lapse sequences, lasting 6 seconds (Fig. 3A). Then, we turn on the flow, to a target *F*_*shear*_, and continue recording images for 36 seconds, during which particles can dissociate. A movie of a typical experiment can be found in Supplementary Video 1. Using a particle tracking algorithm, we quantify the dissociation time, defined as the time between the onset of flow and their removal from their original location, for all of the particles that were present during the baseline phase of the measurement (Fig. 3A). We combine the dissociation times of the 100-400 particles observed in a typical experiment to construct the particle survival probability *p*(*t*), which gives the probability that a particle is still bound after some time *t* after application of the shear force (Fig.3B). Typically, *p*(*t*) takes the form of a stretched exponential function, which is well described by eq. 2.

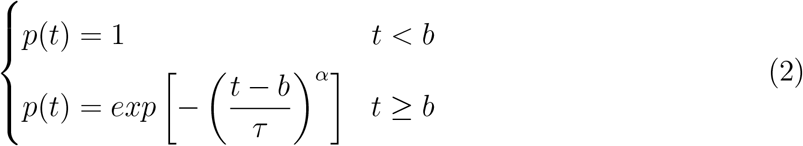

where *τ* is the typical particle lifetime, *α* represents the stretched exponent and *b* denotes an offset that marks the time at which the flow rate is turned on (6 s). Eq. 2 decays more rapidly at higher forces, indicative of a bond stability that decreases with *F*_*shear*_.

**Figure 3:**
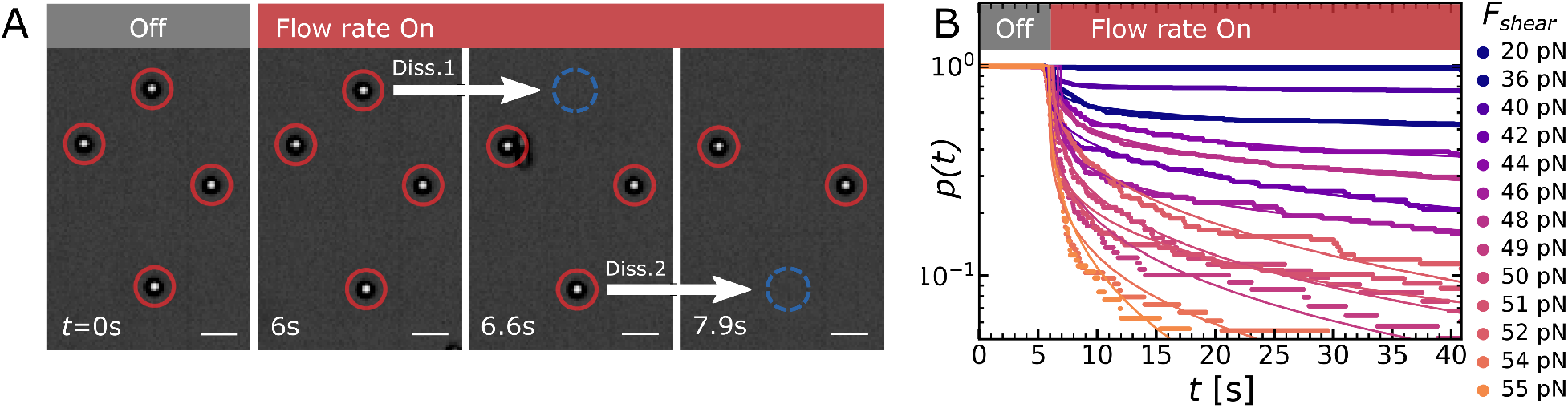
A: Particle dissociation times are determined using a particle tracking algorithm. B: By combining the dissociation times of many particles, we obtain the fraction of bound particles over time, *p*(*t*) which decays more rapidly at varying *F*_*shear*_. Scale bars denote 10 *µ*m.

The observation that our *p*(*t*) curves are well described by a stretched exponential curve suggests that a distribution of characteristic dissociation times is present, rather than a single one, which would give a simple exponential form for *p*(*t*). This distribution of dissociation times is likely the result of the distribution of valencies between the particles and the surface, as was described above in our fluorescence assays of the functionalization. Fitting *p*(*t*) to Eq. 2 allows us to obtain the most probable dissociation time *τ*, which plotted as a function of the applied *F*_*shear*_, results in the force-lifetime curve (Fig. 4A-B).

**Figure 4:**
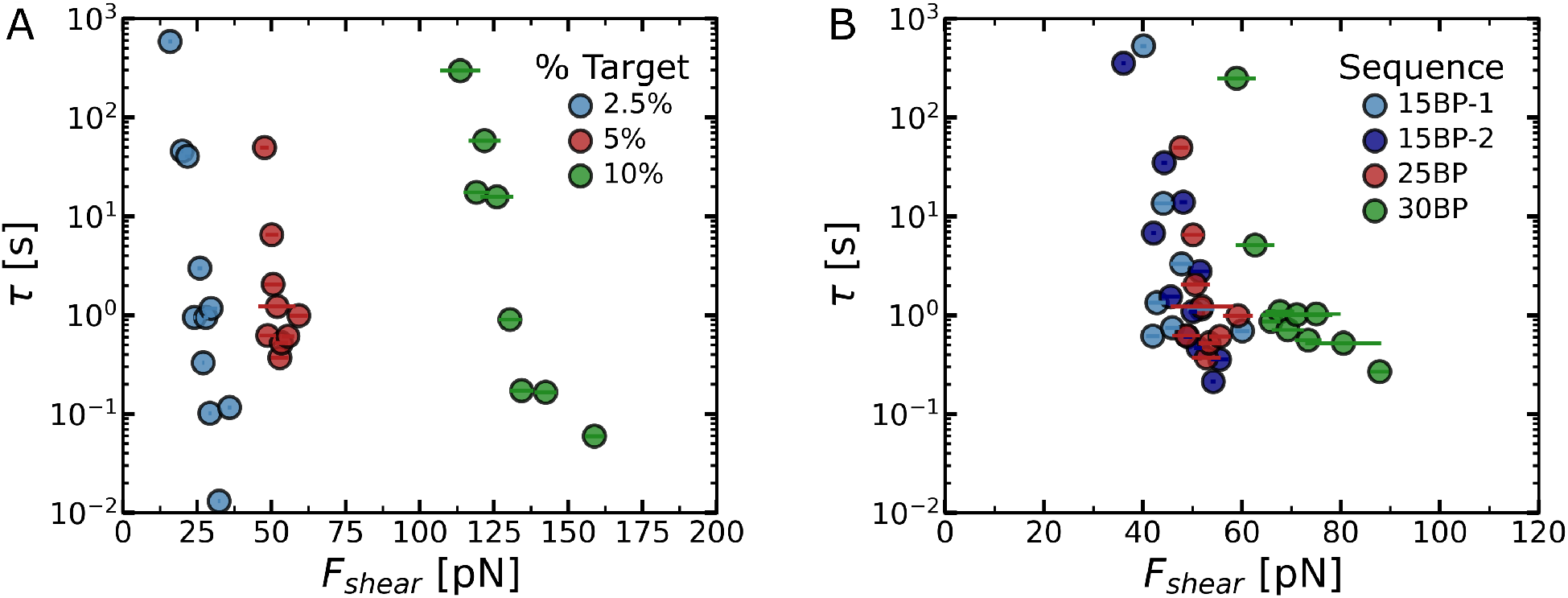
Experimentally determined force-lifetime curves obtained by fitting the *p*(*t*) curves at varying *F*_*shear*_. Two experimental series were performed at (A) varying % target DNA for the 25 bp duplex, and (B) varying DNA sequence lengths at 5 % target DNA. The force-lifetime curve of the 15 bp duplex was recorded twice (15bp-1 and 15bp-2). The DNA sequences are shown in table 2.

As we measure multiple bound particles simultaneously, there is a risk that hydrodynamic coupling between particles affects the applied *F*_*shear*_. Previously performed multiparticle adhesive dynamics simulations on rolling particles have found that hydrodynamic coupling persists on distances up to 5 times the particle radius in the direction of the shear flow, and half that distance in the direction perpendicular to the shear flow. ^23^ To test if hydrodynamic coupling plays a role in our measurements, we have computed the distance of each particle to its nearest neighbouring particle (SI, Section S2). We find that in a typical experiment of 293 bound particles, 89% of particles are at least 5 particle radii removed from their nearestneighbours, and the remaining particles are between 3.3 and 5 radii removed. Therefore, hydrodynamic coupling is unlikely to play a dominant role in our experiments, although it could affect a minority of particles.

Now that we can extract force-lifetime data from the *µ*FFs experiments, we can investigate which molecular properties affect the force-lifetime behavior we observe. We anticipate that the force-lifetime behaviour is affected by both the mechanochemical properties of the bonds and the number of bonds in the particle-surface interface. This is because the *µ*FFS method measures a multivalent interface in which we can expect the shear force to be spread over a large number of bonds simultaneously. If we aim to use the *µ*FFS method as a mechanochemical screening tool, it is essential to know the extent of both these influences. To this end, we have performed *µ*FFS experiments on a series of varying surface densities for a 25 bp DNA duplex, and a series of varying DNA sequence lengths at a constant surface density of 5 % target DNA. The force-lifetime curves obtained in these experiments are shown in figures 4A and 4B, respectively.

First, we investigate the series at varying % target DNA (Fig. 4A). Typically, we observe the particle life time *τ* decays exponentially at increasing *F*_*shear*_. This exponentially decreasing curve is in line with the single-bond Bell-Evans behaviour (Eq. 1), even our assay measures a multivalent interface. The observed decrease in *τ* is remarkably sharp: especially at low fractions of Target DNA of 2.5 %, the observed *τ* can decrease by more than an order-of-magnitude with a change as small as 5 pN in shear force. As the fraction of target DNA is increased between 2.5 % and 10 %, we observe a clear shift of the Force-lifetime curves towards greater shear forces. Increasing the percentage target DNA further resulted in firmly adhering particles that no longer dissociated at the shear forces accessible in this geometry. This trend highlights that the *µ*FFS assay is very sensitive to the bonding density and suggests that users should tune the density of bonds in the particle-surface interface to shift the *F*_*shear*_ - *τ* curves into an experimentally obtainable window.

We proceed to investigate the sensitivity of the *µ*FFS method towards varying the single-bond mechanochemical properties. To this end, we measured the *F*_*shear*_ - *τ* curves of a series of DNA duplexes of lengths 15, 25 and 30 bps at 5 % target DNA (Fig. 4B). We again observe exponentially decaying force-lifetime curves, which shift towards greater *F*_*shear*_ forces as the DNA duplex length is increased. This is explained by the accompanying increase in the mechanochemical strength for these duplexes (Table 1), as established previously by single-molecule force spectroscopy measurements by Strunz et al. ^21^ Upon increasing the duplex length, *τ*_0_ was found to increase by orders of magnitude, which was accompanied by an weak increase in the activation length Δ*x*. We checked the reproducibility of these measurements by performing a technical duplicate of the 15 bp series (15 bp-1 and 15 bp-2), which shows a similar force-lifetime behavior with some minor variations (Fig. 4B).

**Table 1:** Overview of the literature Bell-Evans parameters *τ*_0_ and Δ*x* for the three DNA duplexes used in this study, according to Strunz. et al.^21^ The third column shows the bond densities *ρ* required to map the simulations onto the experimental data.

| Sequence | $\tau_0$ [s] | $\Delta x$ [nm] | $\rho$ [#/$\mu\text{m}^2$] |
| --- | --- | --- | --- |
| 15 bp | $3.2 \cdot 10^4$ | 1.8 | 900 |
| 25 bp | $3.2 \cdot 10^9$ | 2.5 | 600 |
| 30 bp | $1.0 \cdot 10^{12}$ | 2.8 | 900 |

**Table 2:**
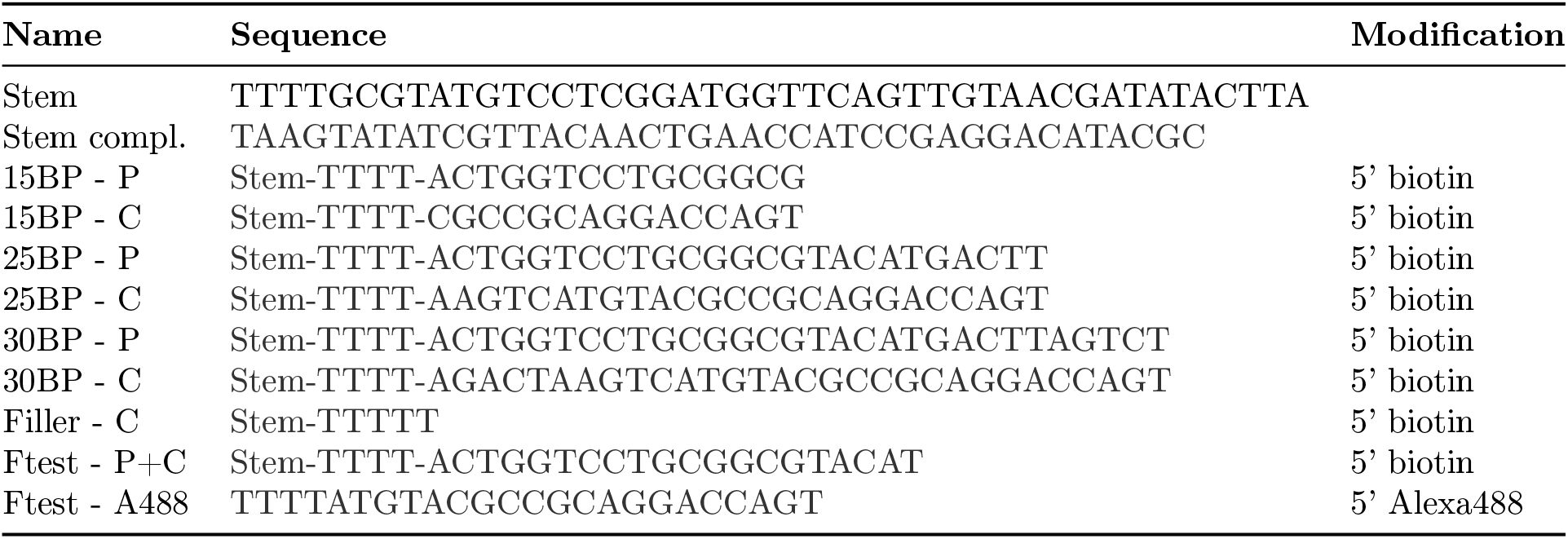
Overview of the DNA oligonucleotides used in this study. Sequences are named according to the length of their sticky ends and whether they were placed on the particle (P) or channel surface (C)

Based on the qualitative observations of the *F*_*shear*_ - *τ* curves, we can conclude that both the bond density and the single-bond mechanochemical properties affect the position of the force-lifetime curves. However, in order to use *µ*FFS as a quantitative mechanotyping method, we require an approach to quantitatively describe these force-lifetime curves in terms of single-bond mechanical properties and bond densities. To this end, we have developed a model system based on quasi-static kinetic Monte Carlo simulations, ^24^ that can be used to simulate the force-lifetime curves using the single-bond *τ*_0_ and Δ*x* and the bond density *ρ* as input parameters (Table 1). Using this model, we can simulate the behavior of a particle bound to the surface as follows (Fig. 5A): 1. First, we place a virtual particle on top of a surface functionalized with linkers at a well-defined density. We allow a fraction of these linkers to form bonds with the particles according to a statistically weighted probability function. 2. We then apply a shear force parallel to the surface of the channel, which applies a shear force and torque on the particle. Using a gradient descent algorithm, we move and rotate the particle until this force and torque is balanced against the forces and torques applied by the bonds and the system achieves mechanical equilibrium. 3. Using a Gillespie kinetic Monte Carlo algorithm,^24^ we randomly allow one bond to break or form, where we take into account the Force-dependent rate constants based on the single-bond mechanical properties, and update the simulation time accordingly. 4. We repeat steps 2 and 3 to simulate how the system behaves over time, until dissociation occurs or a maximum number of steps has passed. By repeating these simulations over a range of *F*_*shear*_ and recording the dissociation times, we obtain simulated force-lifetime curves. Each of the simulations is repeated 20 times to obtain statistics on the lifetime *τ*. A detailed overview of the simulation protocol, the assumptions used, and a motivation of the model parameters can be found in the Supporting Information (Section S1). In these simulations, we can use all-known system parameters and literature values for all the model input parameters, with the exception of the bond formation rate *k*_*on*_, which we estimate at 10^6^ s^*−*1^, and the bond density on the channel surface *ρ*, which we use as the only mapping parameter in our model.

**Figure 5:**
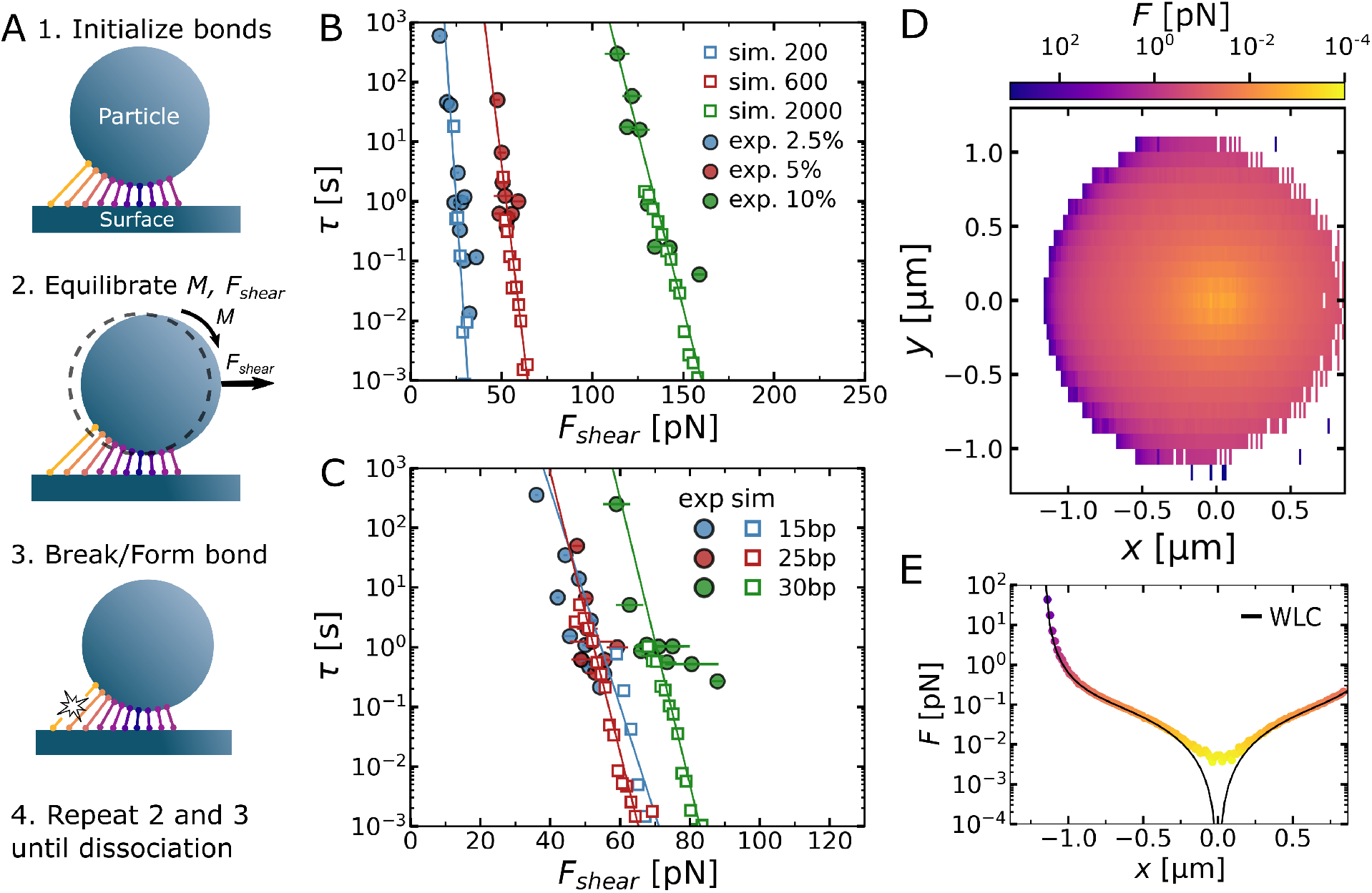
A: Schematic overview of the kinetic Monte Carlo simulations used to simulate the *µ*FFS experiments. B: Simulated force-lifetime curves for he 25 BP duplex at varying surface densities (squares) mapped on the experimental *µ*FFS data (circles). C: Simulated force-lifetime curves for the varying DNA duplexes (squares) mapped on experimental *µ*FFS data (circles). Lines in B and C denote a fit of an exponential decay function through the simulated results. D: Logarithmic heat map (top-down) of the force-distribution on the bonds at the particle-surface interface, averaged over 10 simulations at. E: cross-section taken through the center of the heatmap in C, along with a Worm-like chain curve plotted in black (SI, section S1).

We first performed a simulation series to map the series of measurements at varing % target DNA, by varying the simulated bond surface density *ρ* (Fig. 5B). In these simulations, the single-bond mechanochemical properties of 25 bp DNA as recorded previously in smFS experiments were used as input parameters^21^ (see Table 1). Our simulations produce similar exponentially decaying force-lifetime curves as observed in our experiments, and we obtain an excellent quantitative agreement between the experimental and simulated data. Our simulations further reproduce both the shift of the force-lifetime curves to greater *F*_*shear*_ and the change in slope as the bonding density is increased. This suggests that our modeling approach captures the essential features of the mechanotyping method. We find that the *ρ* values required to map the data at 2.5 %, 5% and 10% Target DNA are 200, 600, and 2000 bonds*/µ*m^2^, respectively. These *ρ* values scale nonlinearly with the applied % target DNA, which is in agreement with our experimental observations in the surface fluorescence tests (Fig. 2B).

Using a similar series of simulations, we map the experimental data at varying DNA duplex lengths (Fig. 5C), using the single-bond Bell-Evans parameters described by Strunz et al.^21^ (Table 1). We again find a good agreement between the simulated and experimental data. This demonstrates that the experimental *µFFS* results can be quantitatively described using the single-bond mechanochemical parameters. As each of these measurements were performed at 5% target DNA, we would expect that the same bond density *ρ* would be required to map these datasets. Interestingly however, we find some variation, with 900 bonds*/µ*m^2^ being required to map the 15 bp and 30 bp data, whereas 600 bonds*/µ*m^2^ is required to map the 25 bp data (Table 1). this could be explained by some variation in the control over the bond density. Taken together, the agreements between our modelling and experiments show that it is indeed possible to describe the particle force-lifetime curves from our method using single-molecule mechanochemical properties, and only the bond density *ρ* as a mapping factor. This opens the way for using the combination of rapid *µ*FFS experiments and simulations as a screening tool to predict single-bond properties from these multivalent measurements.

Besides linking the force-lifetime behaviour to single-bond mechanical properties, our simulations can further provide insight into the dissociation mechanism of the particle-surface interface. This is possible by tracking the force-distribution across the bonds in the interface. We computed the average force in each position relative to the center of the particle, and obtain a heat map of the force distribution from a top-down perspective (Fig. 5D). Strikingly, we find a clear localisation of force on a small number of bonds in a ring at the trailing end of the particle, which is orders of magnitudes higher than the minimal force at the particle center. A cross-section taken through the center of this heatmap reveals that forces of up to 50 pN are reached at the trailing end (Fig. 5E). As the force-lifetime behavior of DNA duplexes is dictated by the Bell-Evans model (Eq. 1), in which the bond lifetime shortens exponentially by force, we can expect this trailing edge to be the main location where bond rupture takes place. This localisation could help explain why the force-lifetime curves measured by the *µ*FFS method are so strongly force-dependent. Due to the strong force localisation, one can expect that as soon as a threshold force is reached that is sufficient to break the few trailing bonds, the particle as a whole will rapidly dissociate.

## Conclusion

In this work, we have presented microfluidic force spectroscopy *µ*FFS as a rapid screening technique for mechanotyping of chemical bonds. We have shown that it is possible to obtain a particle force-lifetime curve in a single experiment, using readily available equipment, and in a much shorter time than typically possible with single-molecule force spectroscopy techniques. Unlike the existing techniques, the *µ*FFS method does not allow one to obtain single-molecule force-lifetime information directly, but rather measures multi-valent interfaces. However, we are able to quantitatively interpret the particle force-lifetime curves in terms of single-molecule mechanochemical properties, by mapping the experimental force-lifetime curves with quasi-static kinetic Monte Carlo simulations. This opens the way to combine the *µ*FFS measurements and simulations into a screening technique for rapid estimation of single bond mechanical properties. As we have found that the quantitative mapping relies strongly on the bond density *ρ* at the channel surface, we expect that a precise control over *ρ* and a method to quantify *ρ* will be essential in future efforts to extract single-bond mechanical properties from these multi-bond *µ*FFS measurements.

## Materials and Methods

All chemicals used in this study were purchased from Sigma Alrich, unless stated otherwise. All DNA oligonucleotides used were ordered from Integrated DNA Technologies (IDT).

### Overview of buffer solutions

The following buffer solutions were used in this work:

- **I**: 10 mM Tris, 1 mM ethylenediaminetetraacetic acid (EDTA), 2 mM MgCl_2_ at pH 6.
- **II**: Buffer I, containing an additional 0.01% w/v Bovine Serum Albumin (BSA) and 0.2 % w/v Pluronic F108
- **III**: Buffer II, containing an additional 100 mM NaCl and 0.2 mM biotin

### DNA sequences and hybridisation protocol

An overview of the DNA sequences and modifications used in this study can be found in table 2. Each oligonucleotide used to functionalise the particle or channel surfaces consists of a 5’ biotin group, followed by 40 bp double-stranded stem, a 4T spacer and finally a sticky end of 15, 25 or 30 nucleotides. The filler DNA lacks this sticky end. This 40 BP double stranded stem was included as a stiff spacer to suppress non-specific interactions between the proteins on both surfaces. First, the double-stranded stem was formed by hybridizing the biotinylated oligonucleotides to the stem complement sequence, using 10 *µ*M of biotinylated oligonucleotide (e.g. 15BP-C) mixed with 10 *µ*M stem complement in buffer I. The sequences were hybridized by incubation for 5 minutes at 95 °C, followed by cooling down to room temperature at a rate of 1 °C*/*min. The hybridized oligonucleotides were stored at 4°C and used within 24 h.

### Functionalisation of polystyrene microparticles

Streptavidin-coated polystyrene particles (4.34 *µ*m diameter) were ordered from Spherotech Inc. (Catalog No. SVP-40-5). Prior to functionalisation with DNA oligos, the particles were transferred to buffer II by washing as follows: 100 *µ*L of 0.5 % w/v particle suspension was centrifuged for 2 minutes at 3000 g, the supernatant was carefully removed with a pipette and the particles were resuspended in 1 mL of buffer II. This washing step was repeated for a total of three times. The particles were then centrifuged one more time and functionalised by resuspending in 80 *µ*L of 10 *µ*M hybridized target (15BP - P, 25BP - P, 30BP - P, table 2). The suspensions where gently shaken for 5 minutes at room temperature, after which they were washed three times in buffer III and finally resuspended in 0.5 mL of buffer III. The functionalised particles were stored at 4 °C and used within two days.

### Functionalisation of microfluidic channels

Thin-bottom flow cells of (1500 *µ*m x 50 *µ*m x 40 mm) ordered from Micronit Microtechnologies B.V. (Art. nr. FC_FLC50.3_PACK) were mounted in a Micronit Fluidic Connect Pro chip holder and connected to 1/32” I.D. 1/16” O.D. polypropylene (PP) tubing. Prior to functionalisation, the flow cells were etched by filling them with 1 M NaOH for 5 minutes using a Luer Lock syringe, and then thoroughly rinsed with 4 mL milliQ water. The inlet and outlet tubing was swiped with pH paper to confirm they were at neutral pH. The subsequent functionalisation steps were performed immediately prior to the *µ*FFS experiment, with the chip holder mounted in the *µ*FFS setup. The chip holder is placed on a Nikon Eclipse Ti2 brightfield microscope with a Fastec HiSpec 2G Mono camera. Flow control was performed using an sequential fluid injection setup (Elvesys), consisting of an Elveflow OB1 Mk3+ microfluidic flow controller, a MUX 12-way bidirectional valve, and an MFS3 2-80 *µ*L/min thermal flowrate sensor.

The flow cell was then functionalised on-flow as follows: First, the channel was incubated in biotinylated Bovine Serum Albumin (Thermo Scientific, cat no. 29130) by flushing 0.5 mL of a 1 mg/mL biotinylated BSA solution in buffer I, after which the flowrate was turned off for 5 minutes to incubate. Next, 4 mL of buffer II was rapidly flushed through at 1.5 bar to remove any unbound biotinylated BSA. The channel was then functionalised with streptavidin, by flushing 0.5 mL of a 0.02 mg/mL streptavidin (Thermo Scientific) solution in buffer II at 50 *µ*L/min, followed by another rinse with 4 mL of buffer II. Next, the channel was functionalised using a mixture of target DNA and filler DNA at the desired target DNA percentage with a total DNA duplex concentration of 3.3 *µ*M in buffer II. Specifically, 0.9 mL of this mixture was flushed through the channel, after which the flowrate was turned off for 5 minutes to incubate. Finally, 4 mL of buffer III was flushed through at 1.5 bar to bring the channels to the ionic strength for the *µ*FFs measurements and block any remaining streptavidin sites with biotin. All *µ*FFs experiments were started within one hour after the aforementioned procedure.

### Fluorescence tests on particles and channel surface

Particles: A batch of polystyrene microparticles was first functionalised with 100% Ftest - P+C DNA (table 2), following the polystyrene particle functionalisation protocol described above. Next, the particles were incubated for 5 minutes in 80 *µ*L of 10 *µ*M Alexa488-labelled complement Ftest-A488 (table 2) in buffer II, followed by three centrifugation-washing cycles with buffer III to remove remaining unbound Ftest-A488 DNA. 50 *µ*L particle suspension was pipetted on a glass coverslip, and images were recorded with a Nikon Ti2 Eclipse confocal fluorescence microscope (60x objective, C2 confocor, 488 nm excitation). Microfluidic channels: Three microfluidic channels were functionalised with 0%, 50% and 100% Ftest - P+C DNA (table 2), according to the microfluidic channel surface functionalisation protocol described above. Next, 0.9 mL of 3.3 *µ*M Ftest-A488 DNA in buffer III was flushed through the channel and the flowrate was turned off for 5 minutes to incubate. 4 mL buffer III was flushed through to wash away the remaining unbound Ftest-A488 DNA. Channels were imaged using the same Nikon Ti2 Eclipse confocal microscope, using a 4x objective.

### Performing the *µ*FFS experiments

A *µ*FFS experiment was started by flushing a batch of DNA-functionalised polystyrene particles into the channel. The flow was then turned off for 5 minutes, allowing the particles to sediment and form interactions with the bottom surface of the chip. A window of 895 *µm* x 716 *µm* was imaged in the center of the microfluidics channel, using a Nikon Plan APO Lambda 20x/0.75 objective. A recording with the Fastec camera was started at 40 fps. After recording for 6 seconds, the flow rate was ramped up to the target flow rate within 1 second and the detachment of particles was recorded over the next 35 seconds while simultaneously recording the applied flow rate through the Elveflow SI software. These experiments were repeated over a range of 15 flow rates per sample. The number of particles present in each measurement was aimed at 100 to 400 particles. This ensures sufficient statistics on dissociation events, while limiting hydrodynamic coupling between particles (SI, section S2).

### Determination of shear force *F*_*shear*_

For each measurement, the average flow rate is determined by averaging the flow rate over the first 10 seconds after turning on the flow, as the great majority of dissociation events occur in this time window. For a rectangular channel filled with Newtonian fluid where the height is much larger than the width (*h << w*), the shear stress at the channel wall can be approximated:

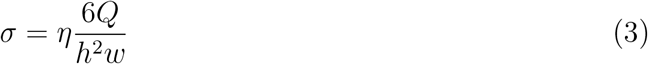

Here *σ* is the shear stress, *Q* is the flow rate, and *η* is the viscosity. The approximation in equation 3 is only valid at the wall in the center of the channel, and breaks down around the edges, where the shear stress deviates to a change in the flow velocity profile. ^25^ In order to apply a well-defined shear stress, it is therefore essential that the areas close to the vertical edges of the channel are avoided in *µ*FFS measurements.We confirmed that *σ* is not position-dependent by plotting a histogram of the dissociation time as a function of particle position (SI, section S6). Since the particles are small compared to the height of the channel, we can assume that the wall shear stress is the shear stress applied across the particle as a whole. This allows us to approximate the resulting force *F*_*shear*_ applied to the particle due to the shear stress using:^19,20^

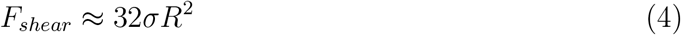

with *R* the particle radius and *σ* the shear stress.

### Particle tracking and data analysis

Particle tracking was performed using a custom-built Python program. The associated code in Python is made publicly available on https://github.com/jorissprakel/uFFS. The following python packages were used: Trackpy, NumPy, SciPy, Matplotlib, Pandas.^26–30^ The raw image data was pre-processed to convert brightfield particle spots into Gaussian spots which could be tracked by the Trackpy package. A detailed description of this process can be found in the supplementary information (section S5). Tracked particles were linked into trajectories allowing for a maximum displacement of 3 pixels between frames, and particles were considered dissociated once they displaced more than 1.4 *µ*m from their resting positions at the start of the experiment, or when the particle trajectory ended. In practice, many trajectories ended immediately after particle dissociation, as the dissociated particles move quickly compared to the framerate.

The particle survival probability (*p*(*t*)) curves were constructed from the particle trajectories by counting the fraction of particles with a dissociation time greater than time *t*. The typical lifetime *τ* is obtained by fitting with a stretched exponential function, as shown in equation 2. Force-lifetime curves were then constructed by plotting *τ* versus the *F*_*shear*_ obtained from flow rate analysis.

### Quasi-static kinetic Monte Carlo simulations

All simulations were performed in a custom-developed python script, making use of the following python packages: NumPy,^27^ and pandas.^30^ The associated code in Python is made publicly available on https://github.com/jorissprakel/uFFS. A detailed description of the simulation protocol, and the assumptions made can be found in the SI, section S1. The simulations in Fig. 5B-C were run for a total of 1 · 10^7^ kinetic Monte Carlo simulation steps. Each simulation was repeated 20 times, and the particle dissociation time *τ* was averaged across these repeats to obtain the data in Fig. 5B-C. To obtain the linker force distributions (Fig. 5D-E), one simulation of the 30 bp duplex, at *ρ* = 900 bonds*/µ*m^2^ was performed for a total of 1 · 10^5^ simulation steps, while the linker position p0 relative to the particle surface and linker extension force F was stored every 10 steps for all linkers. Heat maps were then calculated by dividing the channel surface into a grid of 150 cells along the x-axis and 23 cells along the y-axis, and computing the average extension force *F* of all bonds present within each grid over the course of the simulation.

## Acknowledgement

The work of MvG is financially supported by the VLAG Graduate School. JS and AB are funded by the European Research Council (ERC) project CoG-CATCH. The Authors further thank Lucile Michels for the design of the schematic figures.

## Supporting Information Available

- S1. Description of the quasi-static kinetic Monte Carlo Simulations
- S2. Interparticle separation distances
- S3. Overview of all p(t) curves
- S4. Quantification of DNA binding sites on particles
- S5. Image processing and particle tracking
- S6. Dissociation time distribution along channel width
- Supporting video 1 - video of a typical *µ*FFS experiment.

## TOC Graphic

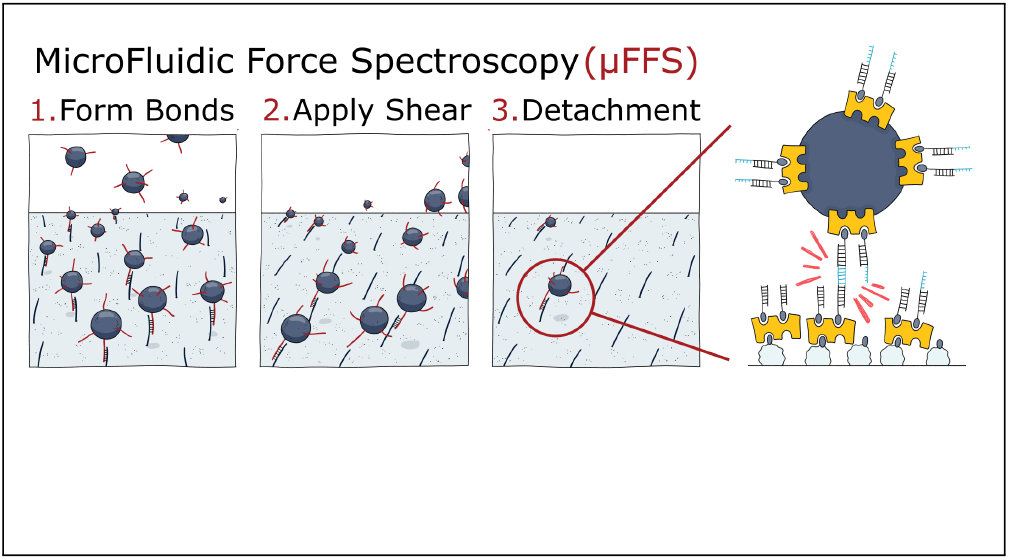

