## Supporting Information for "Rapid Molecular Mechanotyping with Microfluidic Force Spectroscopy"

### S1. Description of the quasi-static Kinetic Monte Carlo Simulations

All simulations were performed in a custom-developed python script, making use of the following python packages: NumPy,[1] and pandas.[2] In this section we describe the simulation protocol in detail. The associated code in Python is made publicly available on https://github.com/jorissprakel/uFFS. An overview of all simulation parameters and the source of the chosen values can be found in the following table:

| **Parameter** | **Description** | **Value** | **Unit** | **Source** |
|---|---|---|---|---|
| $\rho$ | Bond density on channel surface | Variable | $\#/\mu m^2$ | N/A |
| a | Particle radius | 2.45 | $\mu$m | Spherotech inc. |
| $L_{max}$ | Bond contour length | 65 | nm | Calculated[3,4] |
| $L_K$ | persistence length | 0.1 | $\mu$m | Manning, G.S.[5] |
| $k_{on,0}$ | Base bond formation rate | 10ˆ6 | s-1 | Estimate |
| $k_{off,0}$ | Base bond dissociation rate | based on seq. | s-1 | Strunz et al.[6] |
| $\Delta x$ | Activation length | based on seq. | nm | Strunz et al.[6] |

#### Initialization

A simulation is initiated as follows: A simulated particle of diameter 4.34 $\mu$m is placed in a three-dimensional space with its center at a height $h = 2.17$ $\mu$m above a flat plane representing the channel surface, so that it is exactly touching this plane. Next, a rectangular array of binding locations $p_0$ is generated on the surface of the flat plane, such that it obeys the channel surface bond density $\rho$ [bonds/$\mu$m$^2$]. A small amount of random noise on the order of 0.1 times the average distance between binding locations is applied to include a degree of randomness in the system. A number of initial bonds between the binding locations $p_0$ and the particle are then formed using the following protocol:

1. The distance $L$ between each position $p_0$ and the closest point on the particle surface is computed, to determine the length of a potential bond to be formed. Here, we assume

that the bond density on the particle is greater than the bond density on the channel surface, so that a binding location $p_0$ on the channel surface will always find a binding location $p_1$ on the closest position on the particle surface.

2. We compute the extensional free energy change $\Delta G$ [$\mathrm{k_B T}$] according to the Worm-Like-Chain (WLC) model, which has been found to be an accurate model to describe extension of double-stranded DNA duplexes.[7,8]

$$\Delta G = \frac{2k_B T}{L_K}\left(\frac{1}{4}\frac{x^2}{1-x} + \frac{1}{2}x^2 - \frac{0.8}{3.15}x^{3.15}\right) \tag{1}$$

Here, $L_K$ denotes the persistence length and $x{=}L/L_{max}$ is the extension of the duplex relative to its contour length $L_{max}$. The contour length was estimated at 65 nm by adding the sizes of all components of the linkers together, including 2 BSA proteins ($2 \cdot 5.5$ nm)[4] , 2 streptavidin proteins ($2 \cdot 5$ nm)[3] , and 128 nts of DNA, including the two stems, the target sequence, and the spacer T's ($128 \cdot$ 0.34 nm). Chains for which $x > 0.99$ are excluded, as these are stretched to an extent that the chain backbone becomes stretched, so that the standard WLC model becomes invalid. Any bonds formed in this regime would experience so much force that they would immediately dissociate. To keep the simulations running stably, we have excluded these chains.

3. Using a pseudo-random number generator, we then connect bonds randomly such that detailed balance is obeyed:

$$K = \frac{k_{on,0}}{k_{off,0}} \cdot \exp[-\Delta G] \tag{2}$$

where $k_{on,0}$ and $k_{off,0}$ denote the bond formation and dissociation rate constants in absence of force, respectively. Equation 2 takes into account that linkers that have to stretch more in order to form a bond are less likely to form through the use of the Boltzman weighting $\exp[-\Delta G]$, which contains the entropic stretch penalty required to stretch a linker from its rest length to the required length $L$.

#### Equilibration

After initializing the bonds, we apply a shear force $F_{shear}$ and torque $M_{shear}$ to the particle. These values are computed from the applied shear rate via interpolation between the values calculated in the well-established paper by Goldman et al.,[9] table 1. We also compute the extension force applied by each existing bond using the WLC model,[10] as:

$$F = \frac{\delta(\Delta G)}{\delta x} = \frac{2k_B T}{L_K} \left( \frac{1}{4} (1-x)^{-2} - \frac{1}{4} + x - 0.8x^{2.15} \right) \tag{3}$$

with $x = L/L_{max}$ again the extension of the bond relative to its contour length. In the event that a bond experiences a relative extension $x > 0.99$, where the WLC model becomes invalid, the extension force is capped at $F(x = 0.989)$, to prevent computational instabilities. We compute the torques $M_y$ along the y-axis that each bond applies to the particle as:

$$My = (z - h) \cdot F_x - x \cdot F_z \tag{4}$$

where $z$ indicates the height of the linker at the connection point on the surface of the particle ($p_1$), $h$ denotes the height of the center of the particle, and $F_x$ and $F_z$ are the horizontal and vertical components of the extension force computed in Eq. 3.

Next, we equilibrate by minimizing the total energy, which entails balancing all the forces and torques acting on the particle through the shear-induced $F_{shear}$ and $M_{shear}$ using a gradient descent approach. During this minimization, the particle is allowed to translate along the x-axis and rotate around the y-axis, which are the directions in which the shear force and torque are applied. For computational efficiency, we assume that translation and rotation in any other direction is negligible, so that this minimization problem can be limited to just two degrees of freedom.

#### Main simulation loop

After equilibrating the forces and torques, the main simulation protocol is started. In each step of this protocol, we use a Gillespie Kinetic Monte Carlo (KMC) algorithm[11] to randomly allow one bond to break or form, taking into account force-dependent rate constants based on the single-bond mechanical properties, and update the simulation time accordingly. Afterwards, we again equilibrate the force and torque, as described in the "Equilibration" section above, followed by another Kinetic Monte Carlo step. This loop is continued until only one bond remains, at which point the particle is considered dissociated and the current time is stored as dissociation time. If this did not occur before $1 \cdot 10^7$ kinetic monte carlo steps had passed, the particle was not considered dissociated.

The Gillespie Kinetic Monte Carlo steps were performed as follows:

1. First, we compute rate constants for every available bond formation/dissociation step, i.e. every existing bond has a dissociation rate $k_{off}$, whereas every unbound linker has a formation rate $k_{on}$. The rate constants $k_{off}$ is computed using the Bell equation[12] :

$$k_{off} = k_{off,0} \cdot \exp\left[\frac{F\Delta x}{k_B T}\right] \tag{5}$$

   where $k_{off,0}$ is the rate constant in absence of Force and $\Delta x$ is the activation length. $k_{off}$ depends on the linker extension force, which is computed through the WLC model (Eq. 3).

2. For every unbound linker, the formation rate constant $k_{on}$ is computed as:

$$k_{on} = k_{on,0} \cdot \exp\left[\frac{(F\Delta x) - \Delta G}{k_B T}\right] \tag{6}$$

   where $k_{on,0}$ is the base bond formation rate constant, and $F$ and $\Delta G$ are the Forces and free energy associated with forming the bond in question. The choice for equation 6 ensures that the equilibrium constant $K = \frac{k_{on}}{k_{off}} = \frac{k_{on,0}}{k_{off,0}} \exp\left[-\Delta G\right]$, which means that for every extension $x$ the equilibrium $K$ is governed by $\Delta G$, in line with thermodynamic expectations. The inclusion of the free energy change $\Delta G$ in equation 6 also accounts for the effect that DNA duplexes that have to stretch farther to form a bond are less likely to do so. $k_{on}$ is set to 0 for any bond that would be stretched beyond it's contour length to prevent computational instabilities.

3. Once the rate constants for all potential formation and dissociation steps have been calculated, one reaction is selected using a pseudo-random number generator. In this selection process, the rate constants are used as weighing factors, such that those reactions with greater rate constants are more likely to take place. The protocol for this algorithm is described in detail by Gillespie et al.[11] As the Gillespie algorithm is event-driven, the time $\Delta t$ passed between each KMC step is variable, and depends on the rate constants, as described in detail by Gillespie et al.[11] By evaluating $\Delta t$ after each KMC step, the evolution of particle-surface interface can be followed in time. Here we make the assumption that the Force and torque equilibration after each KMC step is fast compared to the bond formation and dissociation rates, such that the relaxation time due to viscous dissipation can be neglected. We expect this assumption is valid, as there are at most times many bonds connected between the particle and the surface, so that each individual bond broken or formed contributes relatively little to the overall force-torque balance.

#### Data storage and further analysis

During the main simulation loop, the simulation time $t$, the particle position $x_P$, and the rotation angle $\theta$ were stored every $1 \cdot 10^4$ simulation steps. The simulations in Fig. 5B-C were run for a total of $1e7$ simulation steps. Each simulation was repeated 20 times, and the particle dissociation time $\tau$ was averaged across these repeats to obtain the data in Fig. 5B-C. To obtain the linker force distributions (Fig. 5D-E), one simulation of the 25 bp duplex, at $\rho = 2000$ bonds/$\mu$m$^2$ was repeated for a total of $1e5$ simulation steps, while the linker position $p0$ relative to the particle surface and linker extension force $F$ was stored every 10 steps for all linkers. Heat maps were then calculated by dividing the channel surface into a grid of 205 cells along the x-axis and 23 cells along the y-axis, and computing the average $F$ of all bonds present within each grid over the course of the simulation. The WLC-curve (black line) in Fig. 5E was obtained by computing the extension length of a bond formed between surface position $x$ and the nearest point on the particle surface using Pythagoras' theorem: $L = \sqrt{x^2 + h^2} - a$, and computing the extension force using Equation 3.

#### S2. Interparticle separation distances

As the $\mu$FFS measurements are performed on a field of view containing a large number of particles in order to obtain good statistics. As a result, there is a risk of hydrodynamic coupling between the bound particles, which could affect the shear forces particles experience. Previously performed multiparticle adhesive dynamics simulations have shown that hydrodynamic coupling between particles plays a clear role at interparticle distances of up to 5 particle radii.[13] We analyzed whether such hydrodynamic coupling could play a role in our experiments, by monitoring the center-to-center distance between each particle and its closest neighbouring particle (Fig. 1A). A histogram of these nearest-neighbour distance taken from a typical $\mu$FFS measurement with 293 particles (25 bp, 5% target DNA) revealed that 81% of particles further than 5 particle radii separated, and the remaining 19 % are between 3.3 and 5 radii separated from their nearest neighbours (Fig. 1B). We therefore expect hydrodynamic does not play a major role on the, although it might affect a minority of the particles.

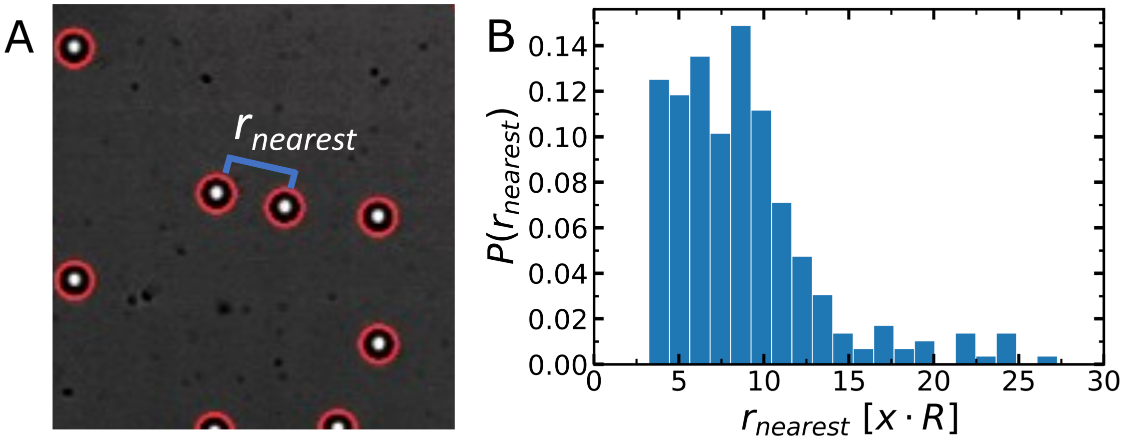


Figure 1: A: The nearest-neighbour distance $r_{nearest}$ is defined as the center-to-center distance between a particle and the closest other particle, expressed in terms of the radius of the particle. B: A histogram of the nearest-neighbor distance taken from a typical measurement with 292 particles.

### S3. Overview of all p(t) curves

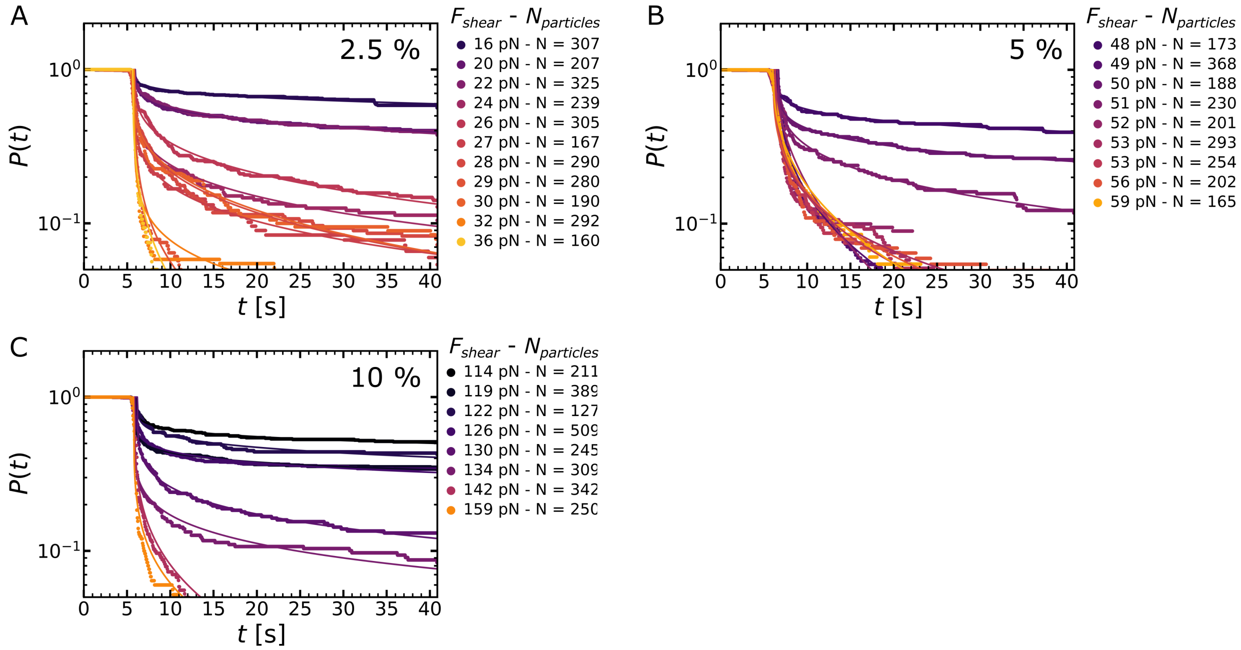


Figure 2: Overview of all $p(t)$-curves recorded for the dataset with varying % target DNA, containing (A) 2.5%, (B) 5%, (C) 10%

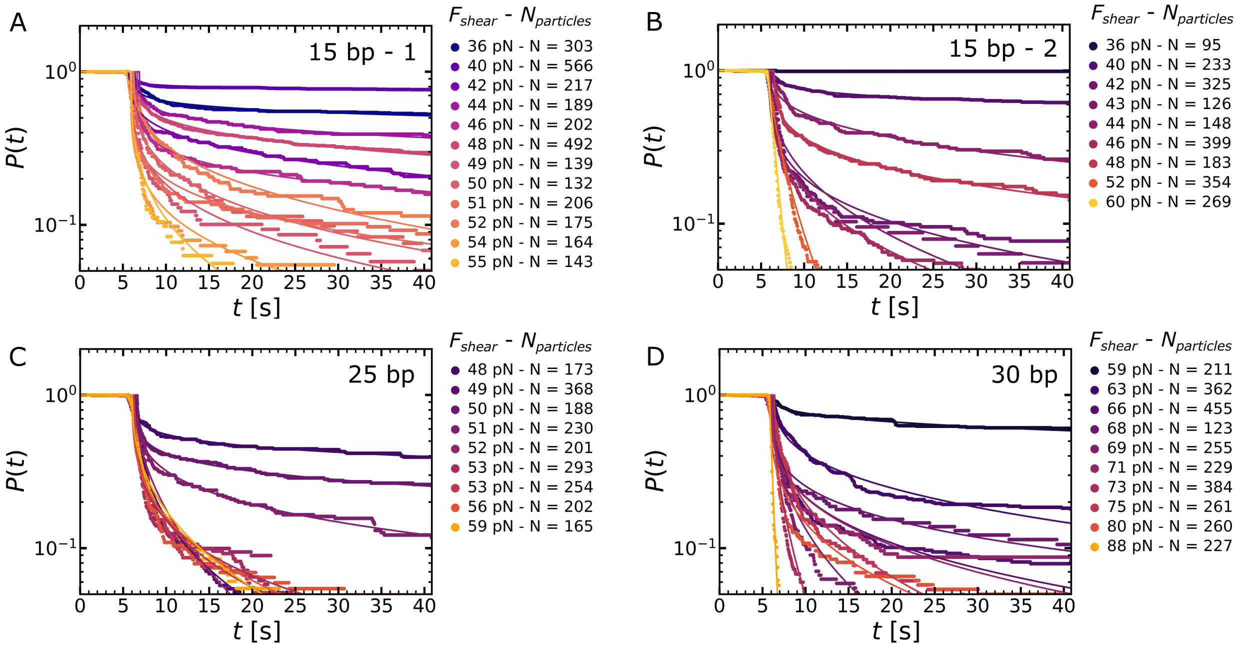


Figure 3: Overview of all $p(t)$-curves recorded for the dataset with varying duplex length at 5% Target DNA, containing (A) 15 bp, first repetition, (B) 15 bp, second repetition, (C) 25 bp, (D) 30 bp. The 25 bp measurement is the same measurement as used in the % Target DNA series in the previous figure.

### S4. Quantification of DNA binding sites on particles

One of the assumptions in our computational model is that the density of DNA binding sites available for binding on the microparticles is substantially greater than the density of available binding sites on the channel surface. To quantify the number of available sites per particle, we have performed a so-called fluorescence pull-down assay, as previously described by Crocker et al.[14] This assay allows one to compute the density of available DNA sites by and measuring the concentration decrease of a fluorescently labelled complementary sequence after incubation with the particles. A batch of 200 $\mu$L 0.5% w/v streptavidin-coated polystyrene particles, was first functionalised with the Ftest - P+C sequence, according to the particle functionalisation protocol described in the Materials and Methods section. Next, the particle suspension is centrifuged for 2 minutes at 3000 xG, the supernatant was carefully removed with a pipette, and the particles were incubated for 5 minutes in an estimated mild excess of 60 $\mu$L 0.22 $\mu$M Ftest - A488 DNA, the fluorescent complement of the Ftest - P+C sequence. After incubation, the particles were again centrifuged for 2 minutes at 3000 G, and the supernatant was carefully collected with a pipette. We quantified the concentration of the Ftest - A488 DNA before incubation and the supernatant after incubation with the particles using a fluorimeter (Fig. 4), and found that 13 pmol of fluorescent DNA had bound to the particles. Using the particle diameter of 4.34 $\mu$m, we computed that this equates to a binding site density $3.9 \cdot 10^3$ sites/$\mu$m$^2$ or a typical distance of 16 nm between each site. The experiment was performed in duplo and yielded similar results. This density is substantially greater than the density of binding sites on the channel $\rho$ we required to map the simulations onto the experimental data (between 200 and 2000 bonds/$\mu$m$^2$). We can therefore conclude that the underlying assumption that the density of DNA sites available on the particle is greater than the density of sites on the channel surface is valid.

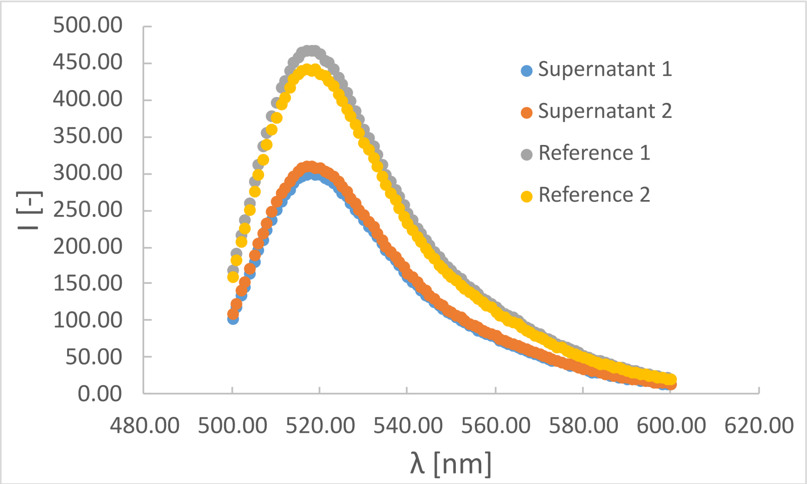


Figure 4: Fluorescence emission spectra of the Ftest - A488 DNA solution before (Reference 1 and 2), and the supernatant after (Supernatant 1 and 2) incubation with the DNA-functionalised polystyrene microparticles show a clear decrease in fluorescence intensity.

### S5. Image processing and particle tracking

The TrackPy particle tracking algorithm used in this study localizes Gaussian features.[15] However, particles localised by brightfield imaging do not resemble Gaussian spots, but rather show as bright spots surrounded by a darker ring (Fig. 5A). Through a series of processing steps, we converted these brightfield features into Gaussian spots that could be accurately tracked with Trackpy. First, the dark ring around each particle (Fig. 5A) was removed using background thresholding, in which any value lower than the most intense pixel in the background was set to zero, followed by a rescaling of the data (Fig. 5B). This yielded a truncated image of the brightfield spots in the center of the particle. This thresholding also ensures that any out-of-focus particles flowing past the bound particles are not tracked, as these have intensities lower than the background threshold. A Gaussian filter of width 1 pixel was then used to convert the truncated images into Gaussian spots (Fig. 5C). Particles were tracked using Trackpy (Fig. 5D). The following TrackPy settings were used in the process: diameter= 9, minmass= 1, maxsize= 6. After tracking, Particle trajectories were linked together using the TrackPy link function, with settings: memory= 9, distance= 3.

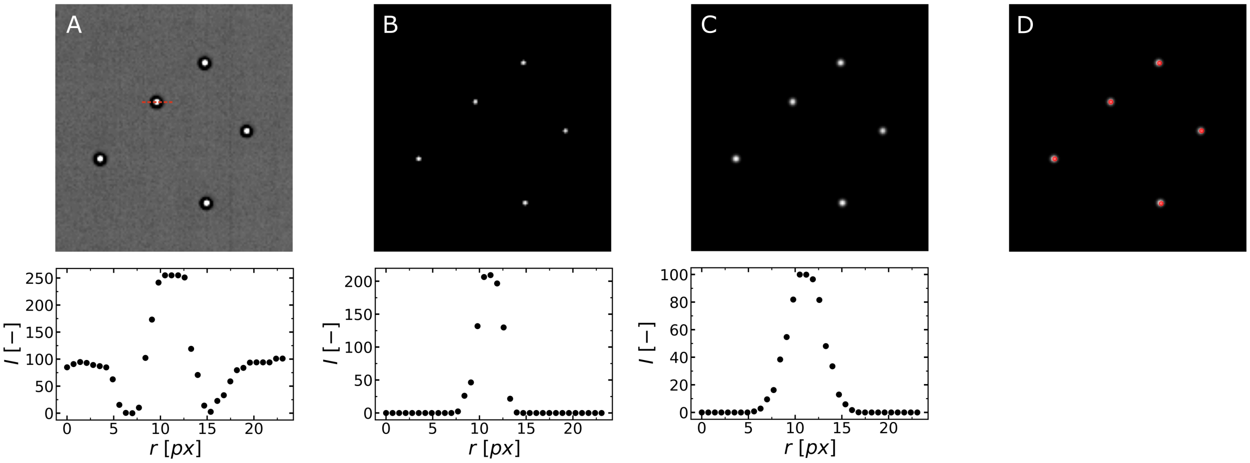


Figure 5: A-D: Overview of the data pre-processing steps prior to particle tracking, showing (top) the image data after each processing step and (bottom) an intensity profile across a single-particle at each step. The following steps were taken: (A) raw images, (B) background thresholding, (C) Gaussian blur, (D) particle localisation. Red dots in (D) correspond to the center of mass of localised particles.

### S6. Dissociation time distribution along channel width

When analysing our $\mu$FFS experiments, we obtain the shear stress $\sigma$ applied to the particles using $\sigma = \eta \frac{6Q}{h^2 w}$. This equation is only valid in the center of the channel, and breaks down at the edges, where $\sigma$ decreases due to a change in flow velocity profile.[16] As it is vital for the $\mu$FFS measurements that same $\sigma$ is applied to all particles in the field-of-view, we have performed all our $\mu$FFS measurements in the center of our channels, in an area of width 895 $\mu$m, as compared to the channel width of 1500 $\mu$m. To confirm that the same $\sigma$ is applied to all particles along the width of this field-of-view, we plotted the distribution of the average particle dissociation times $t_{D,avg}$ over the position $x$ along the width of the channel (Fig. 6). If $\sigma$ decreases along the edge, one expects $t_{D,avg}$ to increase near the edges. This increase is not observed for all measurements except the measurement at $F_{shear} = 48$ , which confirms that there is indeed no spatial variation in $\sigma$. The measurement at $F_{shear} = 48$ does show an increase along one of the edges, although the data is substantially more noisy. We suspect this is an artifact of a lack of statistics, as only 104 particles dissociated over the course of this measurement, corresponding to only 9 particles per $x$-bin.

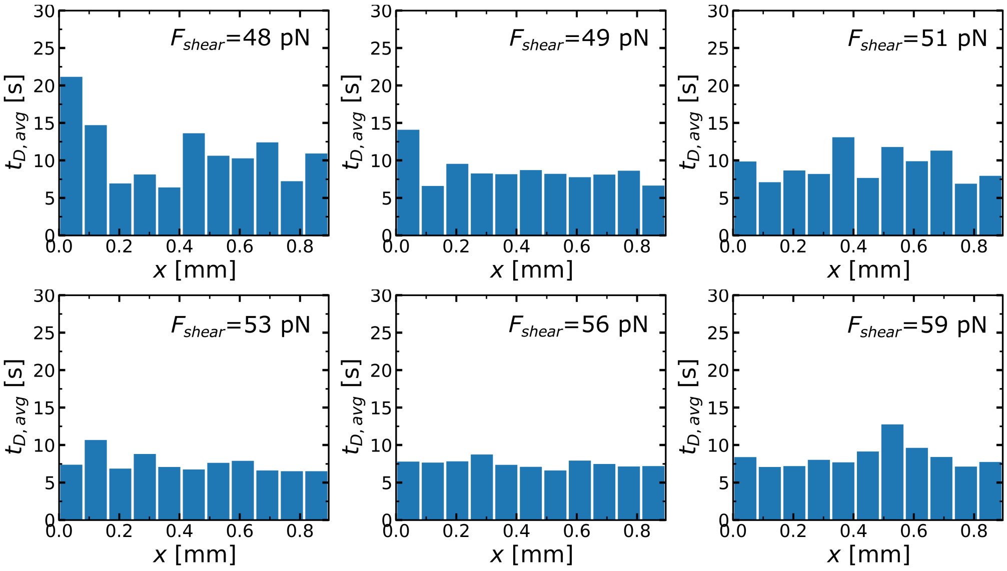


Figure 6: Plots of the average dissociation time $t_{D,avg}$ as a function of the position $x$ along the width of the experimental field-of-view, at varying $F_{shear}$. Data was obtained from $\mu$FFS measurements performed on the 25 bp duplex at 5% target DNA.
